# Extracellular mycobacterial DNA drives disease progression by triggering Caspase-11-dependent pyroptosis of infected macrophages

**DOI:** 10.1101/514125

**Authors:** Monica Varela, Michiel van der Vaart, Arwin Groenewoud, Annemarie H. Meijer

## Abstract

Deregulated inflammation seriously complicates life-threatening microbial infections, including tuberculosis (TB). Assembly of multiprotein inflammasome complexes is an important trigger of inflammation, but how this impacts TB progression remains unknown. Here, *in vivo* imaging in the zebrafish TB model revealed that mycobacterial expansion in TB granulomas is driven by inflammasomes and ensuing pyroptotic cell death of infected macrophages. We show that an Asc-independent pathway induces macrophage pyroptosis and impairs host resistance, in contrast to host-protective roles of Asc-dependent inflammasome activation and Il1b secretion. Using ASC-deficient murine macrophages, we demonstrate extracellular bacterial DNA to induce CASP11-dependent pyroptosis in a manner dependent on phagosome permeabilization. Finally, we propose that mycobacteria induce pyroptosis to escape cell-in-cell structures, formed within granulomas when living infected cells are engulfed by neighbor cells. This study provides new insight into the role of pyroptosis in TB pathogenesis and reveals a novel link between nucleic acid sensing and CASP11-dependent pyroptosis.

## INTRODUCTION

*Mycobacterium tuberculosis* (*Mtb*), the causative agent of tuberculosis (TB), has developed many strategies to manipulate the host immune response (1, 2). The successful establishment of infection depends on the early interactions between pathogen and host innate immune cells, especially macrophages (3). Macrophages play a central role in TB pathogenesis and are the principal niche for intracellular growth of *Mtb* or closely related mycobacterial pathogens, including *Mycobacterium marinum* (*Mmar*), which causes TB in cold-blooded animals (4).

Granulomas, considered as the pathological hallmark of TB, are inflammatory cell structures, rich in macrophages. Granuloma formation starts with aggregation of mycobacterium-infected macrophages and subsequent recruitment of uninfected macrophages. These early stages of granuloma formation favor spreading of bacteria from cell to cell and are dependent on the bacterial RD1 (ESX1) virulence locus (5). At later stages, T-cells and other immune cells are attracted, which are essential to contain the infection. Therefore, granulomas have a dual role, on the one hand serving a host-protective function, but on the other hand enabling dissemination and long-term survival of the pathogen inside its host. Because the failure of a single granuloma to control infection can already be sufficient to initiate disease progression (6), a better understanding of granuloma function is of crucial importance. Many studies have shown that the local balance of pro- and anti-inflammatory responses is critical for determining the fate of individual granulomas and the final disease outcome (7, 8). However, the molecular mechanisms that drive inflammation in the granuloma remain to be elucidated.

The inflammasomes are multiprotein signaling platforms that allow the activation of inflammatory caspases (caspase-1, -4, -5, -11) in response to infections and endogenous danger signals (9-12). Activation of cytosolic pattern recognition receptors that are contained within canonical inflammasomes promotes activation of caspase-1 (CASP1) and interleukin-1β (IL1β) maturation (13). In contrast, the non-canonical inflammasome is dependent on murine Caspase-11 (CASP11) or human Caspase-4/5 (CASP4/5) (14). In response to several Gram-negative pathogens, CASP11 mediates mouse macrophage death via direct interaction with LPS (15, 16). Both inflammasome types activate gasdermin D (GSDMD), which is the executioner of pyroptotic cell death by forming pores in cell membranes (17, 18). Due to the recognition of its necrotic nature, pyroptosis has recently been redefined as gasdermin-mediated programmed necrotic cell death (19).

Despite the known occurrence of necrotic cell death within TB granulomas (20), the role of pyroptosis in granuloma formation and expansion remains unexplored. The well-established zebrafish TB model has been a very useful tool in the study of the disease as it recapitulates many aspects of human tuberculous granulomas (5, 21-23). Studies in zebrafish have shown that granuloma expansion is driven by cell death and subsequent efferocytosis of infected macrophages and that uncontrolled extracellular proliferation of mycobacteria occurs under experimental conditions that promote necrotic rather than apoptotic forms of macrophage cell death (24-26). New tools recently developed for the study of inflammasomes (27) make zebrafish a suitable model for *in vivo* visualization of inflammasomes activation and investigating the role of pyroptosis during granuloma formation.

Here, we report that different inflammasome pathways are activated in infected macrophages during mycobacterial infection. While a pathway dependent on the adaptor protein ASC promotes resistance of the zebrafish host to TB infection, we find that a CASP11 homologous pathway promotes pyroptotic cell death of infected macrophages and exacerbates infection. Using RAW264.7 macrophages, we subsequently show that extracellular bacterial DNA triggers CASP11 activation and GSDMD-dependent pyroptosis, revealing a novel link between nucleic acid sensing and non-canonical inflammasome activation. Finally, we identify the formation of cell-in-cell structures within developing granulomas and demonstrated that *Mmar* is able to escape these structures by triggering pyroptosis, thereby initiating its dissemination and promoting disease progression.

## RESULTS

### 1. Asc AND Il1b ARE HOST-BENEFICIAL DURING *Mmar* INFECTION IN ZEBRAFISH

The interleukin 1 signaling pathway is known to be critical for immunity against tuberculosis (7, 28, 29). To evaluate the role of this pro-inflammatory cytokine in the zebrafish TB model, we transiently knocked down *il1b* in zebrafish larvae. After 2 days of infection, an increased bacterial burden in knockdown (KD) fish was observed (Fig. 1a and Supplementary Fig. 1a), confirming previous evidence for a host-protective role of Il1b during *Mmar* infection (30). In macrophages, maturation and secretion of IL1β occur after activation of inflammasomes (10). For signaling, most of the known inflammasomes require an adaptor molecule, ASC (reviewed in 31). KD of *asc* increased *Mmar* burden in zebrafish larvae (Fig. 1b and Supplementary Fig. 1b). We next investigated ASC oligomerization, considered a hallmark of inflammasome activation. Formation of ASC specks can be visualized just before pyroptosis occurs (32). *In vivo* imaging of zebrafish granulomas showed macrophage pyroptosis occurring after the formation of Asc specks (Fig. 1c). Typically, only one speck per granuloma was observed at a time (Supplementary Fig. 1c). In contrast, Asc speck-independent cell death was more often observed in granulomas (Supplementary Fig. 1d). After pyroptosis, as Kuri et al. (27) showed before, Asc specks can be engulfed and degraded by neighboring macrophages (Fig. 1d, Supplementary Fig. 1e and Supplementary Video 1).

**Figure 1.**
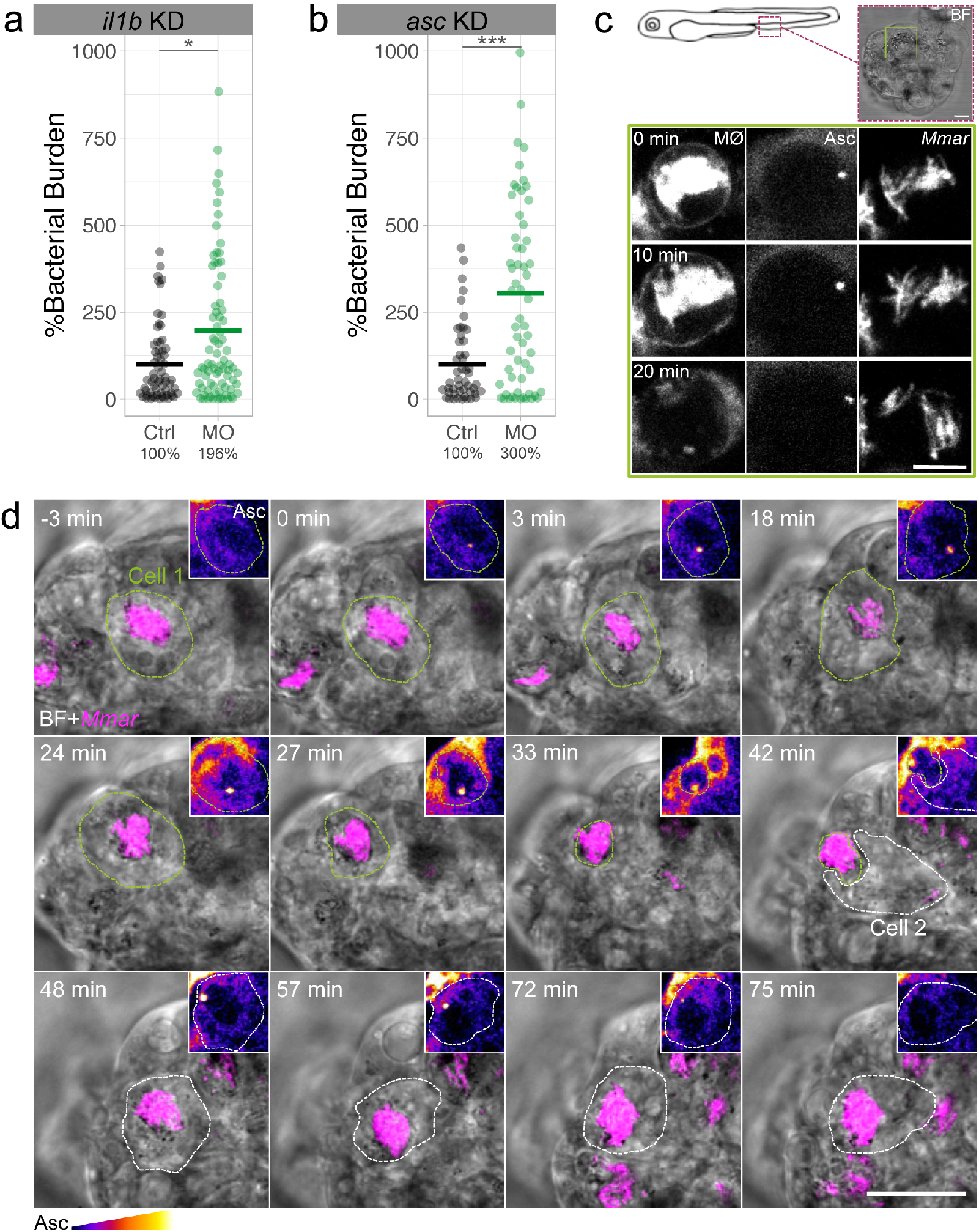
Asc and Il1b are host-beneficial during *Mmar* infection in zebrafish. a, b, Bacterial burden data after transient knockdown (KD) with *il1b* or *asc* morpholinos (MO) at 2 days post-infection (dpi). c, Bright field (BF) and *in vivo* time-lapse confocal images of Asc speck-dependent pyroptosis of a *mpeg1*:mCherry labelled macrophage (MØ) in a zebrafish granuloma at 3 dpi. d, *In vivo* confocal images showing Asc speck formation in Cell 1 (green dotted line) and degradation by Cell 2 (white dotted line) after phagocytosis (see also Supplementary Video 1). Insets show Asc-GFP protein (Fire LUTs). Data is accumulated from 2 independent experiments and representative of at least three independent repeats. Each data point (a, b) represents a single zebrafish larva. Mann-Whitney test (a, b), *p<0.05 and ***p<0.001. Scale bars are 10 (c) and 20 (d) μm.

### 2. TWO INFLAMMASOME SIGNALING AXES ARE DIFFERENTIALLY REGULATED UPON *Mmar* INFECTION

Next, we decided to investigate the role of inflammatory caspases, such as CASP1/4/5/11, indispensable in ASC-dependent and -independent inflammasome signaling pathways (33). Because zebrafish Caspa and Caspb have been traditionally considered CASP1 functional homologues as they can process IL1β *in vitro* (34), we determined the effect of *caspa* and *caspb* transient KD in *Mmar* infected zebrafish. Surprisingly, Caspa and Caspb deficiency resulted in opposite bacterial burden outcomes (Fig. 2a,b, Supplementary Fig. 2a-c). Caspb deficiency increased infection, confirming the results shown by Kenyon et al. (35) using CRISPR/Cas9 to KD *caspb*. In contrast, *caspa* KD or Caspa inhibition with YVAD resulted in a reduced infection burden. Moreover, we found that the double KD of *caspa* with either *caspb* or with *asc* resulted in the same reduced infection phenotype as single KD of *caspa* (Fig. 2c and Supplementary Fig. 2d). In addition, double KD of *caspb* with *asc* had no additive effects in terms of bacterial growth, suggesting that both genes are in the same pathway (Supplementary Fig. 2d). At a cellular level, we observed that while *caspb* KD increased *Mmar* extracellular growth, *caspa* KD was sufficient for restricting the infection (Fig. 2d).

**Figure 2.**
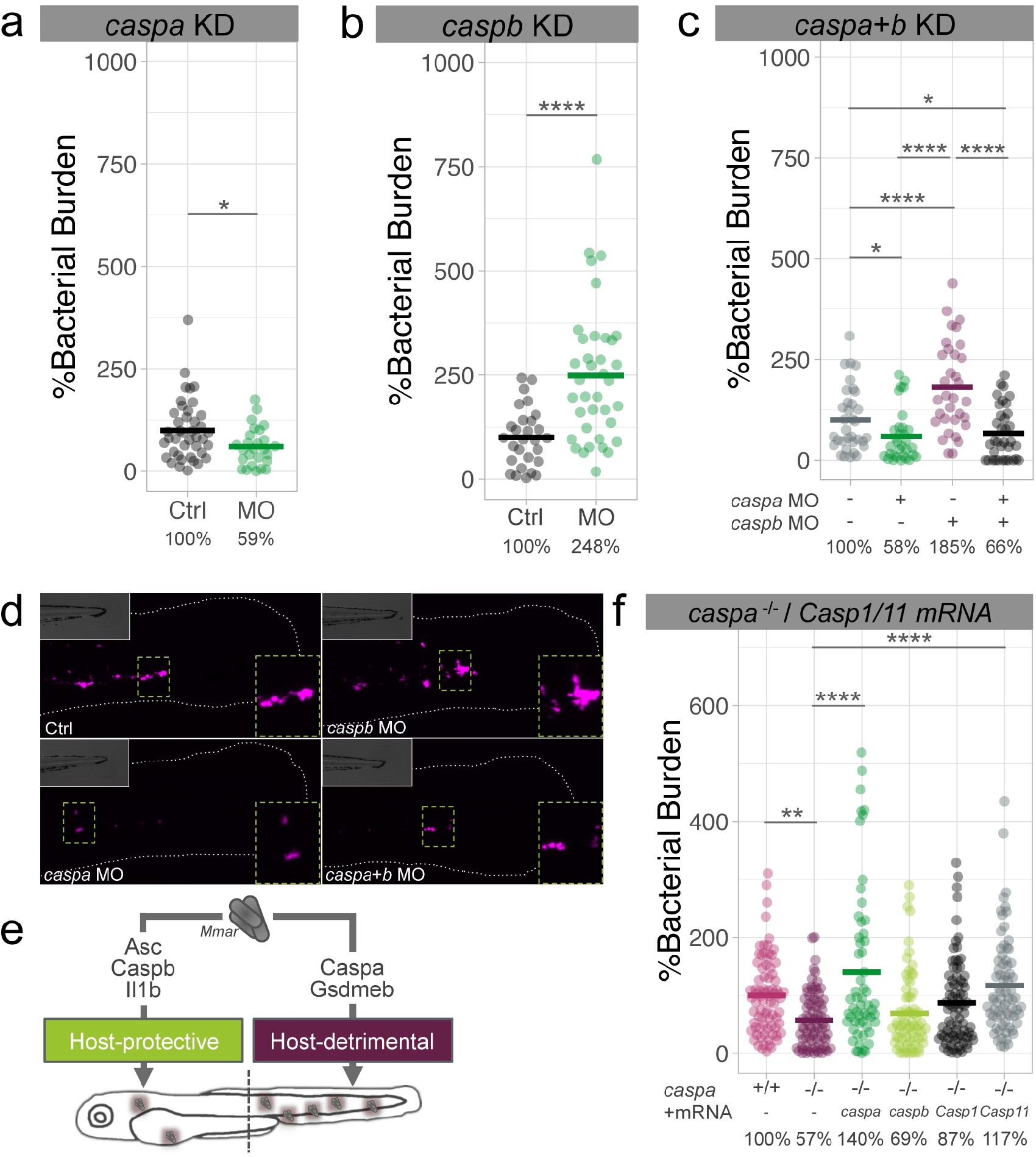
Two differentiated inflammasome axes are activated during *Mmar* infection. a, b, c, Bacterial burden data after transient *caspa*, *caspb* or *caspa*+*caspb* KD in zebrafish larvae at 2 days post-infection (dpi). d, Representative fluorescence images showing differential *Mmar* growth in zebrafish macrophages in the tail fin (outlined and BF images in insets) under different KD situations at 2 dpi. e, Schematic representing the two inflammasome-related axes activated during *Mmar* infection in zebrafish according to our bacterial burden experiments. The model does not exclude other Asc-dependent or -independent roles of Caspa and Caspb under different conditions. f, Bacterial burden data showing the effect of zebrafish *caspa* and *caspb* mRNAs and murine *Casp1* and *Casp11* mRNAs expression in *caspa*^−/−^ zebrafish larvae. Data is accumulated from 2 independent experiments and representative of at least three independent repeats. Mann-Whitney test (a, b) and ordinary one-way ANOVA + Tukey’s multiple comparisons test (c, f), *p<0.05, **p<0.01, ****p<0.0001.

Recently, gasdermin D (GSDMD), which requires processing by inflammatory caspases to form pores in cell membranes, was identified as essential effector of pyroptosis (17, 18). In the zebrafish genome, two proteins with GSDM domain can be identified, Gsdmea and Gsdmeb. *In silico* predictions show that Gsdmeb has a CASP1 cleavage site, which is also the case for GSDMD (Supplementary Fig. 2e). Because of this similarity, we knocked down *gsdmeb* in zebrafish larvae (Supplementary Fig. 2f-g). The lack of Gsdmeb resulted in decreased *Mmar* burden 2 days after infection, similar to *caspa* KD. In addition, the combination of *gsdmeb* KD with *caspa* deficiency had no additive effects (Supplementary Fig. 2h).

Taken together, the results above suggest the involvement of at least two inflammasome pathways during the course of *Mmar* infection, namely a host-protective Asc-Caspb-Il1β axis and a host-detrimental Caspa-Gsdmeb axis (Fig. 2e). This observed duality prompted us to further investigate the functional homology of the zebrafish inflammatory caspases and their murine counterparts. For this, overexpression of zebrafish *caspa* and *caspb*, and murine *Casp1* and *Casp11* mRNAs in *caspa*^−/−^ zebrafish was performed (Fig. 2f). In terms of bacterial burden, only zebrafish *caspa* and murine *Casp11* mRNAs were able to fully rescue the *caspa* mutant phenotype, demonstrating that Casp11 can compensate for the loss-of-function of Caspa and suggesting that in the context of *Mmar* infection the host-detrimental Caspa-Gsdmeb axis in zebrafish is equivalent to the mammalian Casp11-Gsdmd pathway.

### 3. *Mmar* TRIGGERS ASC-INDEPENDENT CELL DEATH AFTER PHAGOSOMAL MEMBRANE RUPTURE

To fully understand the dynamics of inflammasome activation in zebrafish, we explored the formation of ASC specks in the granuloma in combination with *in situ* caspase activity detection, using a fluorescently labeled inhibitory substrate (Flica YVAD) specific for zebrafish Caspa (36). As observed earlier, Asc specks appeared in close contact with bacterial clusters (Fig. 3a). *In situ* staining revealed that active Caspa was abundant in granulomas, visible either in small specks or in patchy patterns around *Mmar* (Fig. 3A, Supplementary Fig. 3a-b). We found that Caspa activation occurred in Asc specks and, in an Asc-independent manner, in both Caspa specks and patches (Fig. 3B and Supplementary Fig. 3c). This, together with our bacterial burden assays, led us to propose that while Caspa can be activated in an Asc-dependent and -independent manner, its Asc-independent activation is the one driving *Mmar* infection progression. We next assessed whether ASC is required for induction of macrophage pyroptosis *in vitro* after *Mmar* infection (Fig. 3c). Using ASC-deficient RAW264.7 macrophages (37) we observed pyroptosis occurring 16 minutes after bacterial phagosome rupture (Fig. 3d and Supplementary Video 2). To confirm the role of phagosomal rupture in the induction of pyroptosis we used mutant *Mmar* lacking the RD1 virulence locus. We found that *Mmar* ΔRD1 could not induce pyroptosis in RAW264.7 macrophages (Fig. 3e). To further demonstrate the importance of phagosomal membrane rupture in the dissemination of infection we compared bacterial burdens in zebrafish after infection with *Mmar* ΔRD1 in *caspa* KD conditions (Fig. 3f). We found that Caspa deficiency had no effect on bacterial burden when bacteria were not able to escape the phagosome, suggesting that phagosomal membrane permeabilization is required for Caspa activation and ensuing pyroptotic cell death.

**Figure 3.**
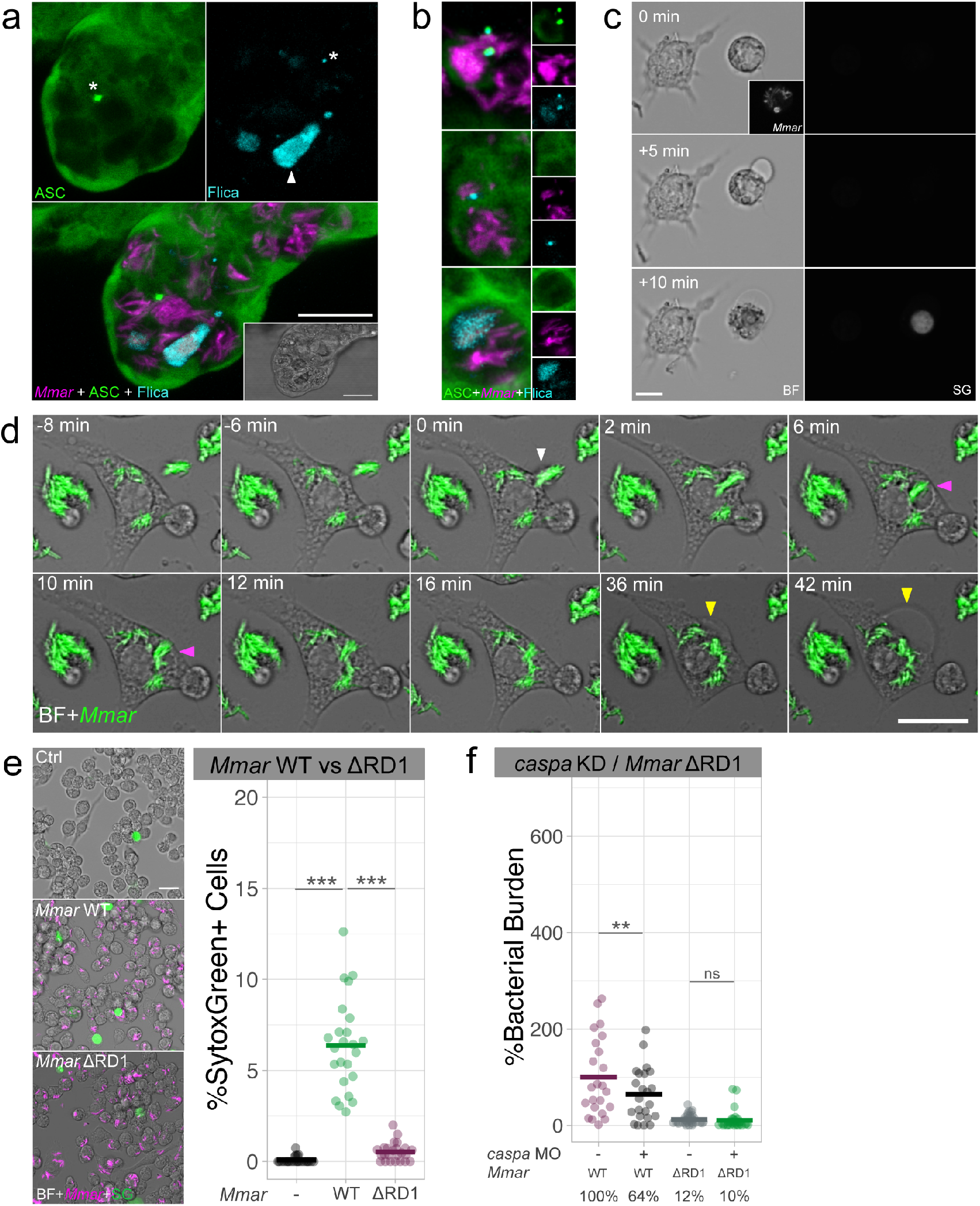
*Mmar* triggers ASC-independent and phagosome rupture-dependent macrophage pyroptosis. a, Confocal images (with BF inset) of a zebrafish granuloma showing Asc and *in situ* Caspa activity (Flica) at 3 days post-infection (dpi). Different types of specks (asterisks) and patches (arrowhead) are observed during *Mmar* infection. b, Representative confocal images showing types of Asc/Caspa activation patterns observed during *Mmar* infections in zebrafish. c, Confocal images showing Sytox Green (SG) uptake in ASC-deficient RAW264.7 macrophages upon *Mmar* infection. d, Confocal footage showing phagocytosis (white arrowhead), phagosome formation and subsequent rupture (magenta arrowheads) and pyroptosis (yellow arrowheads) in RAW264.7 macrophages after *Mmar* infection (see also Supplementary Video 2). e, Sytox Green uptake quantification in RAW264.7 macrophages at 6 hours post-infection (hpi) with *Mmar* WT and ΔRD1. f, Bacterial burden data after transient *caspa* KD in zebrafish larvae 2 days after *Mmar* WT and ΔRD1 infection. Data is representative of three independent experiments. Ordinary one-way ANOVA + Tukey’s multiple comparisons test (e, f), **p<0.01, ***p<0.001. Scale bar are 20 (a) and 10 (c, d, e) μm.

### 4. *Mmar* INFECTION TRIGGERS CASPASE-11 AND GASDERMIN D-DEPENDENT PYROPTOSIS

A previous report showed that CASP1 was not required for *Mtb*-induced cell death in primary mouse macrophages (38). Because our zebrafish data implicated CASP11 in the dissemination of *Mmar* infection, we asked whether or not pyroptosis is CASP11 dependent in murine macrophages. CASP11 cleavage was detected in the supernatant of RAW264.7 macrophages infected with WT *Mmar* (Fig. 4a and Supplementary Fig. 4a). In contrast, CASP11 was not processed upon *Mmar* ΔRD1 infection. In agreement, infection with WT *Mmar* led to high levels of lactate dehydrogenase (LDH) release, while the lack of LDH release after infection with *Mmar* ΔRD1 indicated that cell death was not induced (Fig. 4b). Additionally, knockdown of *Casp11* and *Gsdmd* in RAW264.7 macrophages diminished *Mmar*-induced cell death (Fig. 4c and Supplementary Fig. 4b). Having demonstrated the importance of CASP11 and GSDMD-dependent cell death *in vitro*, we next investigated its role *in vivo*. Using TUNEL staining, which marks apoptotic and pyroptotic cells, we detected reduced numbers of cell death events following *caspa* and *gsdmeb* KD in *Mmar*-infected zebrafish (Fig. 4d and 4e). In conclusion, both *in vitro* and *in vivo* data support the requirement of CASP11/Caspa and GSDMD/Gsdmeb for *Mmar*-dependent pyroptotic cell death.

**Figure 4.**
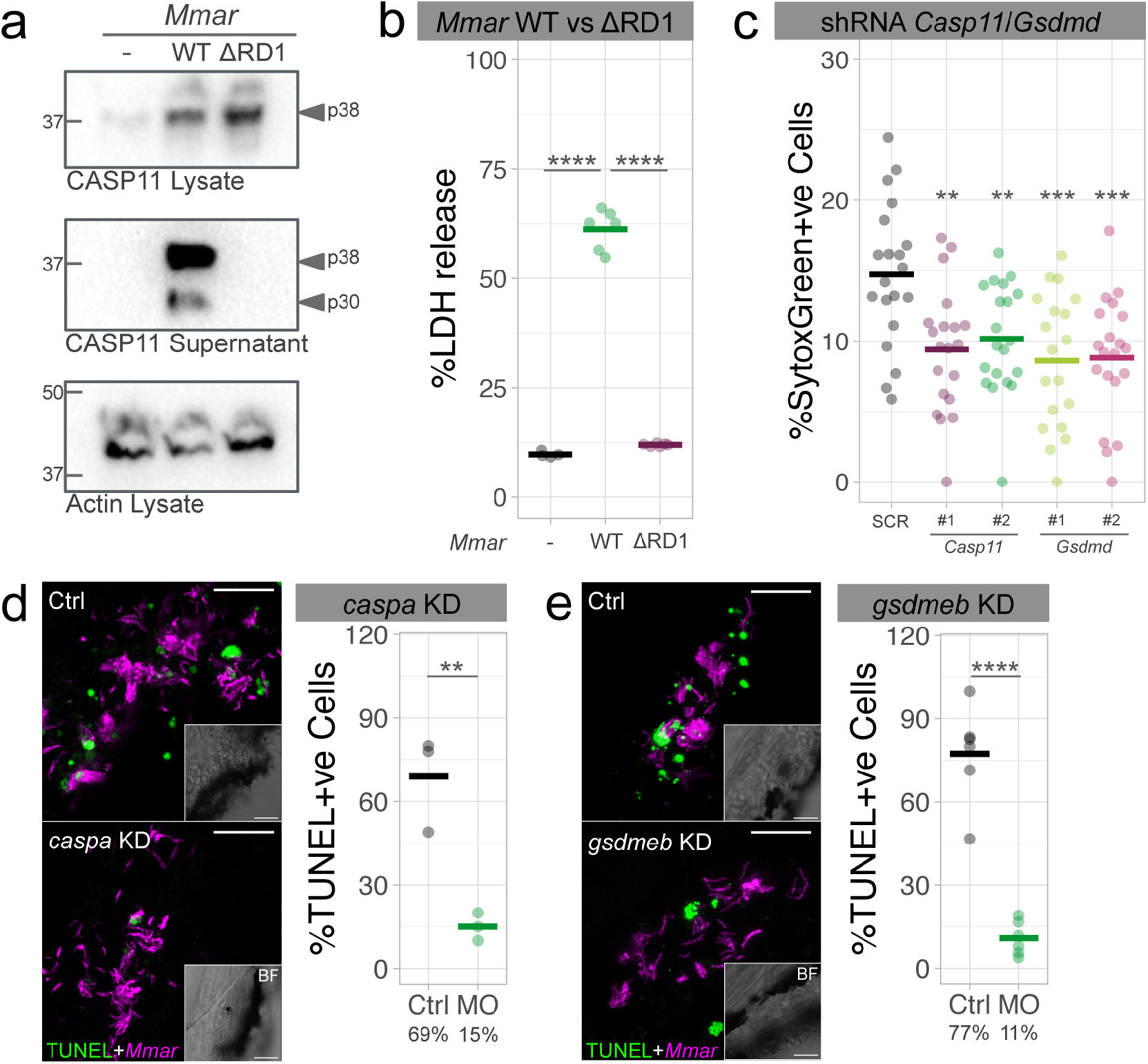
*Mmar*-induced pyroptosis is CASP11/GSDMD-dependent in murine macrophages and Caspa/Gsdmeb-dependent in zebrafish larvae. a, Casp11 protein expression in cell lysates and supernatants of RAW264.7 macrophages after 16 hours of infection with *Mmar* WT and ΔRD1. b, LDH release in RAW264.7 macrophages infected for 16 hours with *Mmar* WT and ΔRD1. c, Cell death (Sytox Green uptake) in scramble control (SCR), *Casp11* KD (x2 different shRNAs) and *Gsdmd* KD (x2 different shRNAs) RAW264.7 macrophages at 6 hpi with *Mmar*. d, e, Quantification of TUNEL-positive *Mmar*-infected cells in Ctrl and *caspa* or *gsdmd* KD zebrafish larvae at 3 days post-infection. Data is representative of three independent experiments. Mann-Whitney test (d, e) and ordinary one-way ANOVA + Tukey’s multiple comparisons test (b, c), **p<0.01, ***p<0.001, ****p<0.0001. Scale bars are 20 (d, e) μm.

### 5. MYCOBACTERIAL EXTRACELLULAR DNA TRIGGERS CASPASE-11 AND GASDERMIN D-DEPENDENT PYROPTOSIS OF INFECTED CELLS

We next sought to identify the bacterial component that induces CASP11 cleavage and GSDMD-dependent pyroptosis of infected macrophages. Unexpectedly, while investigating cell death during infection in zebrafish, we frequently observed positive TUNEL staining other than cell nuclei around *Mmar* (Fig. 5a). To identify the source of this DNA we stained the bacterial culture with the DNA dye Draq5 prior to the infection of RAW264.7 macrophages (Fig. 5b). During infection, we observed Draq5-positive DNA around *Mmar* while the cell nuclei remained negative, indicating the bacterial origin of this DNA. In multicellular bacterial communities of several genera, including *Mycobacterium,* it is common to observe extracellular DNA (eDNA) of high molecular weight, similar to genomic DNA, without signs of massive bacterial lysis (39). To explore the possible involvement of such eDNA in pyroptosis induction, we simulated this situation by coating *Mmar* with ultrapure *E. coli (Ec)* genomic DNA (gDNA) (Fig. 5c), endotoxin-free *Mmar* gDNA, CpG oligonucleotides (ODN2395), and *Ec* LPS as a known CASP11 activator. We found that LPS and both *Mmar* and *Ec* genomic DNAs, but not ODN2395, were able to increase pyroptosis of RAW264.7 macrophages (Fig. 5d). This effect was not observed during infection with *Mmar* ΔRD1, consistent with the inability of this strain to induce phagosome rupture (Fig. 5e). Importantly, we found that cell death induced by bacterial eDNA is CASP11- and GSDMD-dependent (Fig. 5f).

**Figure 5.**
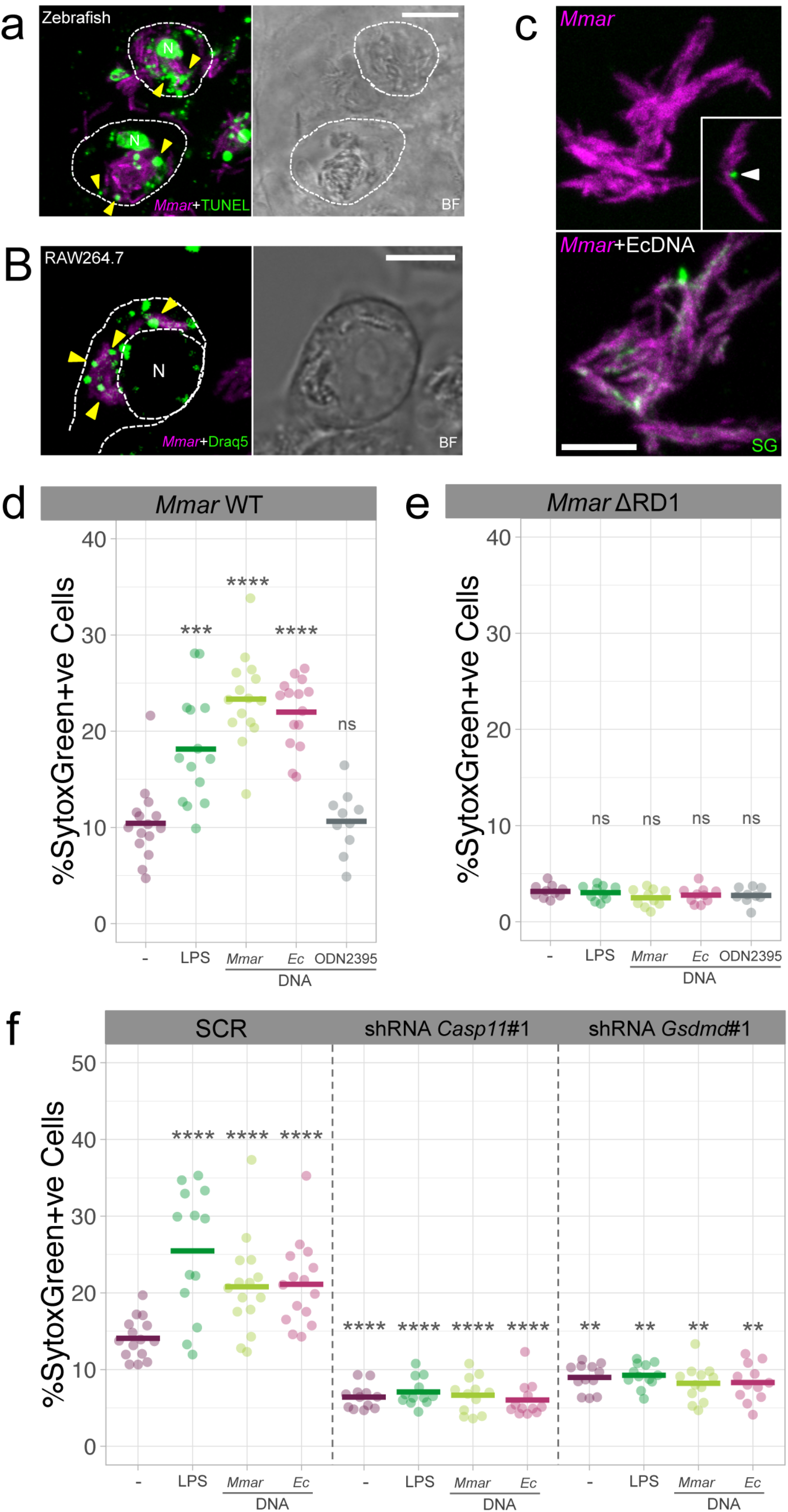
Extracellular *Mmar* DNA triggers CASP11/GSDMD-dependent pyroptosis. a, Confocal images showing TUNEL-stained infected zebrafish cells (outlined) with positive DNA signal in the nucleus (N) and around *Mmar* (arrowheads). b, Confocal images of RAW264.7 macrophages infected with Draq5 (DNA) O/N-stained *Mmar* (MOI 10, 3 hours post-infection (hpi)). N indicates cell nucleus and arrowheads indicate DNA positive staining around *Mmar*. c, Confocal images of a *Mmar* culture incubated with Sytox Green (SG) to stain *Mmar* DNA and a *Mmar* culture incubated with *E. coli* genomic DNA and Sytox Green to visualize the DNA coating. Inset shows an example of *Mmar* with extracellular DNA, occasionally observed in culture. d, Sytox Green uptake quantification in RAW264.7 macrophages infected with *Mmar* WT and *Mmar* WT coated with LPS, *Mmar* genomic DNA, *Ec* genomic DNA or ODN2395 (6 hpi). e, Sytox Green uptake quantification in RAW264.7 macrophages infected with *Mmar* ΔRD1 and *Mmar* ΔRD1 coated with LPS, *Mmar* genomic DNA, *Ec* genomic DNA or ODN2395 (6 hpi). f, Sytox Green uptake quantification in SCR control, *Casp11* and *Gsdmd* shRNA RAW264.7 macrophages infected with *Mmar* WT and *Mmar* WT coated with LPS, *Mmar* genomic DNA, Ec genomic DNA or ODN2395 (6 hpi). Data is representative of three independent experiments. Ordinary one-way ANOVA + Tukey’s multiple comparisons test (d, e, f), **p<0.01, ***p<0.001, ****p<0.0001. Scale bar are 10 (a, b) and 5 (c) μm.

### 6. GRANULOMA FORMATION AND *Mmar* DISSEMINATION *IN VIVO* IS RELATED TO THE BACTERIAL ABILITY TO TRIGGER CASPASE-A ACTIVATION

*In vivo*, proliferation of *Mmar* is related to its ability to escape from phagosomes and induce granuloma formation (Fig. 6a-b). The inability of *Mmar* ΔRD1 to form typical mycobacterial granulomas and trigger Caspa activation (Fig. 6c), suggests that Caspa activation is required for *Mmar* dissemination *in vivo*. Indeed, active Caspa was highly detected in granuloma cells (Fig. 6d).

**Figure 6.**
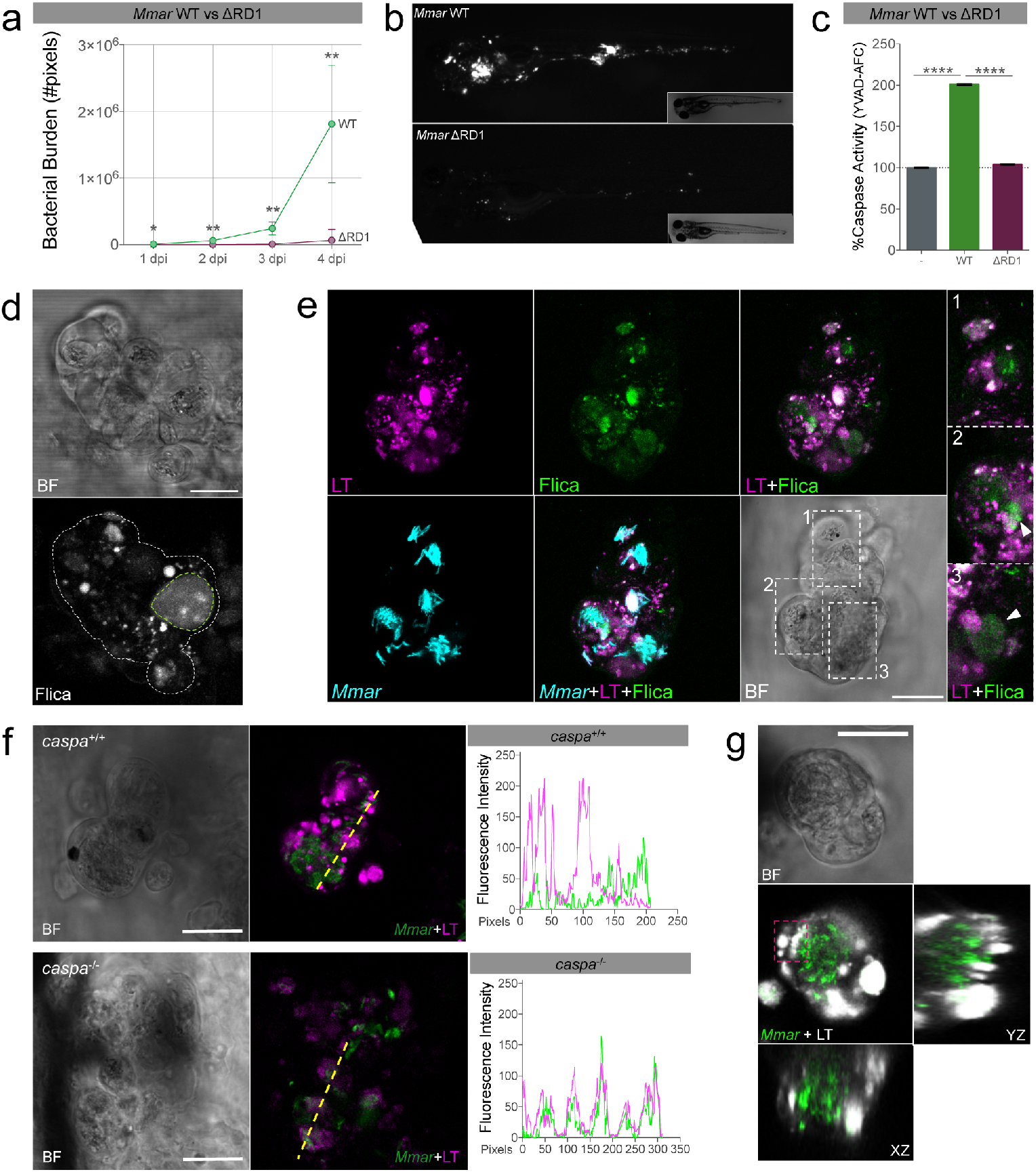
Caspa activation is required for the formation of granulomas in zebrafish. a, Bacterial burden from 1 to 4 days post-infection (dpi) after injection of WT and ΔRD1 *Mmar* in zebrafish embryos. b, Representative fluorescent images at 4 dpi showing the absence of granuloma formation during *Mmar* ΔRD1 infection in zebrafish. c, Caspase activity (YVAD-FMK) in non-infected, *Mmar* WT and *Mmar* ΔRD1-infected larvae 4 dpi (n=2, 40 zebrafish larvae/n). d, Representative confocal images showing *in situ* Caspa activation (Flica) in a zebrafish granuloma at 3 dpi with WT *Mmar*. e, Confocal images of a granuloma stained with LysoTracker (LT) and Flica. Insets 1, 2 and 3 showing Flica-positive and LT-negative phagosomes (arrowheads). f, Confocal images showing the altered acidification pattern (LT) in zebrafish granulomas due to the lack of Caspa. Fluorescence intensities of *Mmar* and LT from the yellow transects are represented on the graphs to visualize the co-localization between both markers. g, Confocal images of LT ring-like patterns around *Mmar* in zebrafish granulomas (square). Orthogonal views are displayed to show the lack of co-localization between *Mmar* and LT. Data is representative of three independent experiments. Mann-Whitney test (a) and Ordinary one-way ANOVA + Tukey’s multiple comparisons test (c), *p<0.05, **p<0.01, ****p<0.0001. Scale bars are 20 (d, e, f, g) μm.

In recent years, inflammasomes have been associated with phagosome acidification and maturation during infection (40). To investigate the possible implication of Caspa in the maturation of *Mmar*-containing phagosomes we analyzed *in situ* Caspa activation in relation to acidification of organelles using LysoTracker (LT) staining. We observed that some bacterial clusters within the granuloma are Flica+ve but LT-ve, which can be interpreted as Caspa activation preceding or preventing phagosome acidification (Fig. 6e). In line with this, we found that the lack of Caspa alters the acidification pattern of bacteria-containing vesicles (Fig. 6f). Acidic vesicles accumulated around *Mmar* but did not colocalize with bacteria in wildtype fish. However, in absence of Caspa, LT staining and *Mmar* fully colocalized (Fig. 6f). These results suggest a non-inflammatory role of Caspa related to vesicular trafficking during *Mmar* infection, as it was previously reported for CASP1/CASP11 during other intracellular bacterial infections (41-43). Moreover, the investigation of *Mmar*-LT colocalization in granulomas revealed the presence of LT patterns concentrically surrounding *Mmar* clusters (Fig. 6g). Because such lysosomal ring patterns are a known cytological feature of entosis (44), this observation suggests the presence of cell-in-cell structures in mycobacterial granulomas.

### 7. *Mmar* ELICITS PYROPTOSIS TO ESCAPE CELL-IN-CELL STRUCTURES IN GRANULOMAS

We next decided to investigate the role of cell-in-cell structures in the fate of the infection. The presence of cell-in-cell structures in granulomas is made visible using cytoplasmic ASC as cell marker that allows visualizing the entotic vacuole devoid of cytoplasmic material between inner and outer cells. While inner cells are always infected, this is not the case for the outer cell that engulfed the infected inner cells (Fig. 7a and Supplementary Fig. 5a). We observed the formation of such multinucleated cell structures *in vivo* (Fig. 7b and Supplementary Fig. 5b) and *in vitro* during the course of *Mma*r infection (Supplementary Fig. 5c). Time-lapse analysis revealed cell-cell contacts occurring before cell invasion (Supplementary Video 3). Moreover, we observed how the outer cell transfers bacteria into an inner cell (Supplementary Fig. 5c and Supplementary Video 3). Taken together, these results suggest an active role for the outer cell, most likely as a consequence of the cell-cell communication happening before engulfment.

**Figure 7.**
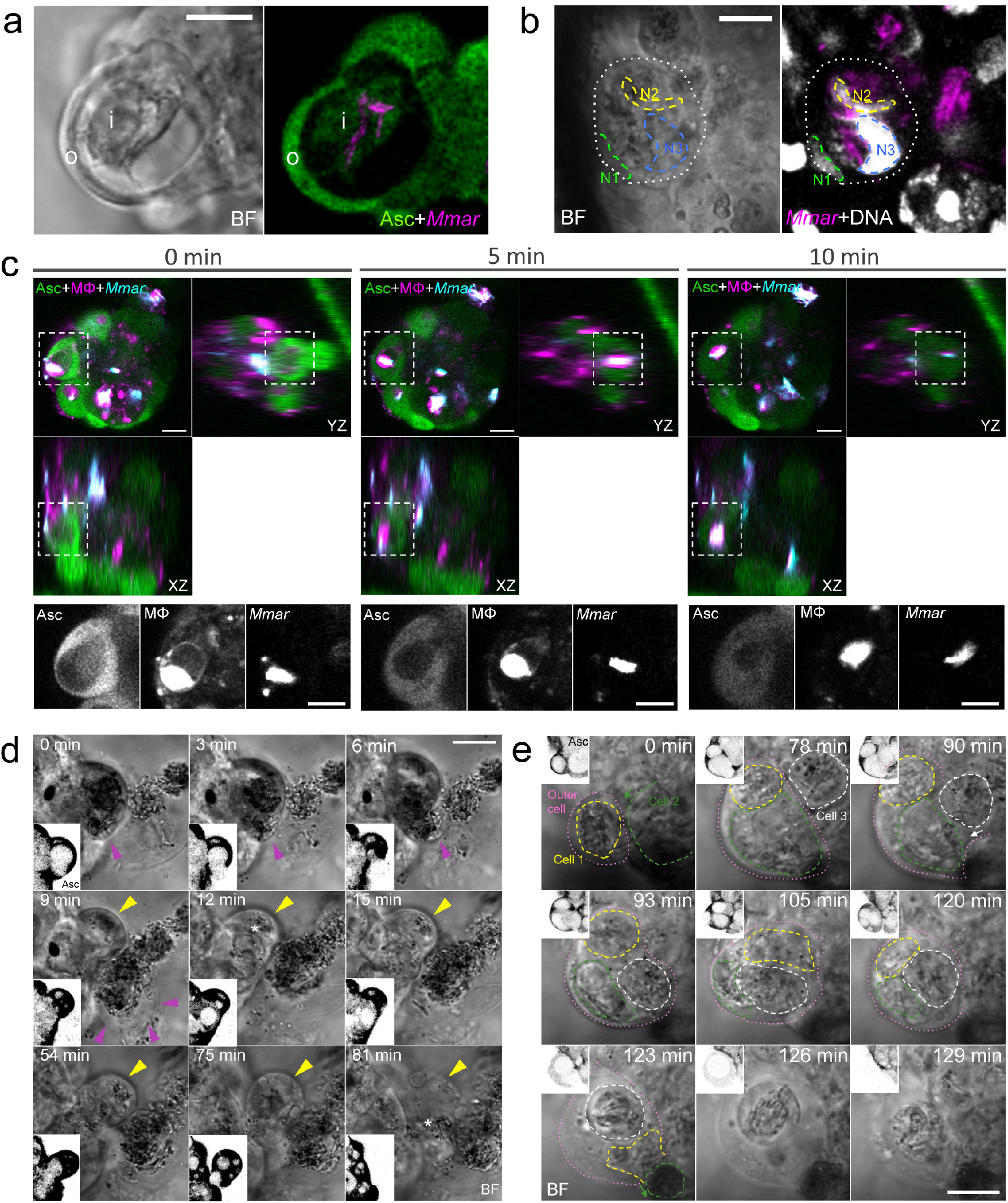
After cell-in-cell structure formation in granulomas, pyroptosis increases *Mmar* dissemination. a, Confocal images of a cell-in-cell structure in a zebrafish granuloma where cytoplasmic ASC is visible in outer (o) and inner (i) cells. b, Confocal images showing a multinucleated cell-in-cell structure in a zebrafish granuloma. The white outline indicates the boundary of the outer cell and the coloured outlines indicate the nuclei (N1-3) of inner and outer cells. Cell 2 (N2) and Cell 3 (N3), both infected, are inside Cell 1 (N1). c, Confocal images showing the death of the inner cell (infected macrophage) in a granuloma cell-in-cell structure. Orthogonal projections are shown to facilitate the visualization of the inner cell killing. d, Confocal images showing the release of an inner cell from the cell-in-cell structure following the initiation of pyroptotic cell death (magenta arrowheads). The infected outer cell (asterisk) eventually dies via pyroptosis (yellow arrowheads) 1 hour later. Insets show Asc (inverted LUTs) for a better visualization of the process (see also Supplementary Video 4). e, Confocal images showing how a single cell in a zebrafish granuloma (pink dotted lines) can uptake more than 1 infected cell (Cell 1 in yellow, Cell 2 in green, Cell 3 in white). Insets show Asc (inverted LUTs) expression. Asc-independent cell death is visible in Cell 3 and in the outer cell (see also Supplementary Video 5). Scale bars are 10 (a, b, c, d, e) μm.

Several mechanisms have been described where the engulfment of whole live cells, leading to the formation of cell-in-cell structures, induces cell death (45). In *Mmar* granulomas we observed engulfment of infected macrophages (Asc+ve, Mpeg1+ve) and their degradation inside outer cells, an epithelial cell in this case (Asc+ve, Mpeg1-ve) (Fig. 7c). Besides a role for cell engulfment mechanisms in killing inner cells, we observed the expulsion of infected inner cells in granulomas (Fig. 7d, Supplementary 5d and Supplementary Video 4). This regurgitation follows the initiation of the pyroptotic cell death program of the inner cell, which in some occasions resulted in pyroptotic dead of both the inner and outer cell (Fig. 7e, Supplementary Fig. 5e and Supplementary Video 5). Importantly, pyroptosis of infected cells was host-detrimental as *Mmar* was subsequently able to proliferate and expand the extracellular infection foci (Supplementary Fig. 6a). Similarly to results obtained by Mahamed et al. (46) during *Mtb* infection, we observed that extracellular *Mmar* attracts new phagocytic cells to the area, which die via pyroptosis shortly after phagocytosis, thereby functioning as a mechanism for rapid expansion of the infection (Supplementary Fig. 6b and Supplementary Video 6). Based on these results, we conclude that pyroptosis of internalized infected cells during entosis in granulomas favors *Mmar* survival and dissemination.

## DISCUSSION

Inflammasome signaling is central to many anti-bacterial host responses. Specifically, CASP11 and its human orthologues CASP4 and CASP5 are emerging as key regulators of this pathway, but their involvement in pathological inflammation during TB remains unexplored. Based on *in vitro* data in murine macrophages and *in vivo* data in zebrafish larvae, we propose CASP11/Caspa as a critical mediator of the dissemination of mycobacterial infection. We found that phagosome permeabilization is required for induction of CASP11/Caspa- and GSDMD/Gsdmeb-dependent pyroptosis in response to bacterial eDNA. Furthermore, we provide evidence that *Mmar* directs the activation of CASP11/Caspa-dependent pyroptosis to escape cell-in-cell structures in granulomas. Together, these findings unravel a mechanism exploited by the bacteria to survive and disseminate within the host.

In agreement with recent reports on macrophages *in vitro* (47, 48), our data suggest that Il1b processing and pyroptosis might be uncoupled during *Mmar* infection in zebrafish. Deregulated production of IL1β worsens TB and therefore it needs to be tightly regulated in the course of the disease (7, 28, 49). In our infection model, IL1b processing seems to occur mainly downstream of an Asc-dependent inflammasome and the host-protective function of this pathway is supported by increased infection burden under both *il1b* and *asc* knockdown conditions. ASC aggregates are stable and can trigger prolonged inflammation even when they are released to the extracellular space (50, 51). In this regard, the host might be able to regulate the number of specks forming to avoid an excessive and detrimental inflammatory response within the granuloma. Due to the highly inflammatory nature of TB in its active state (disseminated tuberculosis), a comparison with sepsis is of interest. Both *Gsdmd*^−/−^ and *Casp11*^−/−^ mice survive after a challenge with lethal doses of LPS (17, 18). In line with this, our data show that CASP11/Caspa and GSDMD/Gsdmeb activation is host-detrimental *in vitro* and *in vivo* during *Mmar* infection. It remains to be established if CASP11 and GSDMD activation have similar disease exacerbating roles during infection with *Mtb*.

We found that macrophage pyroptosis requires phagosome permeabilization, which is directly related to the ability of *Mmar* to activate CASP11/Caspa as *Mmar* lacking the RD1 locus/ESX-1 system cannot activate CASP11/Caspa. ESX-1 is responsible for translocation of eDNA to the cytosol during mycobacterial infections (52, 53). We observed that genomic DNA from *Mmar* and *Ec* can activate pyroptosis *in vitro* in a CASP11-GSDMD-dependent manner to the same magnitude as LPS. CASP11 can bind directly to intracellular LPS, resulting in CASP11 processing and activation (15), but whether a direct interaction between eDNA and CASP11/Caspa also exists will require further research. Recent studies reveal a novel function for guanylate-binding proteins (GBPs) in controlling the intracellular detection of LPS derived from extracellular bacteria in the form of outer membrane vesicles (54). Moreover, in response to certain cytosolic bacteria, GBPs liberate microbial DNA for activation of the DNA-sensing AIM2 inflammasome (reviewed in 55). Intriguingly, the AIM2 inflammasome is an ASC-dependent inflammasome, indicating that CASP11 activation in ASC-deficient RAW264.7 macrophages upon *Mmar* infection must occur via a different pathway. Secretion of eDNA has recently been proposed as a pro-survival strategy in bacterial communities (56). The architecture of these communities depends on the production of an extracellular matrix, of which DNA is part (57). Interestingly, most of the known mechanisms of eDNA release are regulated by quorum sensing, and therefore eDNA is usually produced in response to an increase in the cell density of the population (58). In mycobacteria, stress and quorum sensing result in the formation of biofilm-like structures and persistent cells in necrotic areas (59), suggesting that eDNA secretion is not only host-detrimental because of its ability to induce pyroptosis of infected cells, but also because it can facilitate extracellular proliferation of bacteria.

Notably, we identified cell-in-cell structures in *Mmar* granulomas. Cell-in-cell structures were described decades ago, but their biological significance is only just beginning to be understood. Within the last few years, several forms of cell-in-cell death have been described, among which entosis and cannibalism are of special interest in cancer research and tumor prognosis (60). The main difference between these two processes is the cell taking the lead during engulfment. During cell cannibalism the outer cell leads and engulfs inner cells, while during entosis, it is the inner cell that pushes itself into the outer cell (61). Besides the differences, both processes end in lysosomal digestion of inner cells (62, 63). Regulated by nutrient signaling pathways, entosis is induced under starvation conditions (64). In the context of *Mmar* infection, degradation of inner cells is visible within granulomas, but it remains to be explored whether this mechanism is involved in acquisition of cell nutrients or is primarily a measure of infection containment. One other infection study has previously reported on cell-in-cell structures, revealing that Epstein-Barr virus uses an in-cell infection mechanism to spread from infected B cells to epithelial cells (65).

Granuloma macrophages can undergo multinucleated giant cell (MGC) formation or epithelioid transformation and epithelial cadherin (E-cadherin) has been proposed as a common link between these two differentiation pathways (66). Interestingly, entosis is induced by the expression of E-cadherin proteins (67), pointing to the possible implication of entosis in forming MGCs in granulomas. In zebrafish, macrophage-specific disruption of E-cadherin function resulted in disordered granuloma formation, enhanced immune cell access, decreased bacterial burden, and increased host survival (22). This would suggest that the formation of cell-in-cell structures in granulomas, if dependent on E-cadherin, could benefit the pathogen. The fates of the inner cells and outer cells in cell-in-cell structures are diverse (68). Our data demonstrate that pyroptosis is induced by virulent *Mmar* in inner cells to escape cell-in-cell structures in granulomas, thereby contributing to the expansion of granulomas and infection progression. How pyroptosis induction occurs in the context of cell-in-cell structures will be the focus of future research. Our data suggest that Caspa activation precedes phagosome acidification and therefore corroborate that CASP11/Caspa is involved in vesicular trafficking during *Mmar* infection and it may also be affecting lysosomal degradation and the fate of inner cells. In this regard, the inner cell could be considered like a big phagosome inside the outer cell (a matryoshka doll-like or nested structure). Recently, cleaved GSDMD was found in mitochondria membranes (69) and future research will reveal if this is the case also in phagosome membranes, as GSDMD cleavage by CASP11 might be facilitating bacterial escape from phagosomes, in a similar way as pyroptosis is implicated in the escape of inner cells from outer cells in cell-in-cell structures in granulomas.

Overall, these findings highlight the importance of cell-cell interactions in TB pathogenesis. Local inflammation, the immune response and bacterial state all contribute to the fate of a granuloma. Therefore, there are multiple pathways that can be manipulated to either control infection or promote bacterial dissemination. CASP11/Caspa-dependent pyroptosis increases bacterial survival and dissemination, not only allowing the bacteria to grow in the extracellular space but also increasing infection of new cells. We propose a central role for CASP11/Caspa in regulating vesicular trafficking during infection as well as mediating mycobacterial escape from cell-in-cell structures in granulomas. Based on these results, human CASP4/5 might be a valuable biomarker for monitoring TB therapy and its role during infection should be taken into account for the design of new therapeutic interventions.

## METHODS

### Zebrafish husbandry and infections

Zebrafish (*Danio rerio*) were handled in compliance with the directives of the local animal welfare committee of Leiden University (License number: 10612) and maintained according to the standard guidelines from the Zebrafish Model Organism Database (http://zfin.org). All protocols adhered to the international guidelines specified by the EU Animal Protection Directive 2010/63/EU. All zebrafish experiments were done on embryos or larvae up to 5 days post fertilization (hpf), which have not yet reached the free-feeding stage. Embryos were grown at 28.5°C and kept under anesthesia with egg water containing 0.02% buffered 3-aminobenzoic acid ethyl ester (Tricaine, Sigma) during bacterial injections, imaging and fixation. The following zebrafish lines were used in this study: AB/TL, Tg(*mpeg1.1:mCherryF*)^ump2Tg^, Tg(*asc:asc-EGFP*)^hdb10Tg^ (37) and the mutant line *caspa*^hdb11^ (37).

*Mycobacterium marinum* (*Mmar*), M and ΔRD1 strain, expressing mCherry (70, 71), E2-Crimson (72) or Wasabi (72) fluorescent proteins were grown according established protocols (73). Fresh bacterial inoculums were injected into 30 hpf zebrafish embryos via blood island injection following previously described methods (73). A total of 300 cfu of *Mmar* in a volume of 1nl were injected per embryo in all experiments performed in this study.

### Morpholino oligonucleotides

Morpholino oligonucleotides (MO) were designed and synthesized by Gene Tools (Philomath, OR). Previously validated MOs were used in this work: IL1b i2e3 (0,5 mM): 5’-CCCACAAACTGCAAAATATCAGCTT-3’ (74), ASC atg (0,6 mM): 5’-GCTGCTCCTTGAAAGATTCCGCCAT-3’ (37), CASPa atg (0,6 mM): 5’-GCCATGTTTAGCTCAGGGCGCTGAC-3’ (36). New MOs were validated during this research: GSDMEb e2i2 (0,7 mM): 5’-TCATGCTCATGCTAGTCAGGGAGG-3’, CASPb e2i2 (0,5 mM): 5’-AATCAAAATACTTGCATCGTCTCCG-3’. These MOs were validated by PCR using the following primers: Caspb validation F: 5’-ATGGAGGATATTACCCAGCT-3’, Caspb validation R: 5’-TCACAGTCCAGGAAACAGGT-3’, GSDMEb validation F: 5’-CAAGCGTAACCGATACTGGT-3’, GSDMEb validation R: 5’-AATCCACCTTTGACTGCTGG-3’. Morpholinos were diluted in sterile water with 1% phenol red sodium salt solution (Sigma) and 1 nl of the above mentioned final concentration was injected into the cell of 1 cell-stage embryos.

### Cell Culture and infections

Mouse macrophage-like RAW264.7 cells were maintained in 50% Dulbecco’s modified Eagle’s medium (DMEM) and 50% F12 medium with 10% fetal calf serum, at 32°C under a humidified atmosphere of 94% air and 6% CO2. All infections in this study were performed at a multiplicity of infection of 10.

For WB experiments cells were seeded in 6-well plates (Greiner) at a density of 2·10^6^ cells/well the day before the infection. Cells were infected at a MOI of 10 for 2 hours at 32°C. The extracellular bacteria was washed twice with PBS and removed. Cells were further incubated at 32°C for 14 hours. For cell death assays (Sytox Green uptake) cells were seeded in μ-Slide 8 well glass bottom (IBIDI) at a density of 1·10^5^ cells/well and infections were performed for 6 hours. LDH assay was performed 16 hours after infection using 5·10^4^ cells following manufacturer instructions (Pierce LDH Cytotoxicity Assay Kit, ThermoFisher Scientific).

For coating experiments 100 ul of a 100 cfu/nl bacterial suspension was incubated for 2 hours in 1mg/ml solution of *Ec* O111:B4 LPS (Sigma), *Mmar* M gDNA, *Ec* K12 double stranded gDNA (InvivoGen), or ODN2395 (InvivoGen) previously cell infection. After profuse washing of bacterial suspensions, cells were infected. A bacterial suspension sample was taken to perform a staining using SytoxGreen and visualize the coating efficiency. *Mmar* gDNA was isolated and endotoxin-cleaned using established protocols (75, 76).

### mRNA synthesis

For *caspa*, *caspb*, *Casp1* and *Casp11* retrotranscription we used mMessage mMachine T7 Ultra (ThermoFisher Scientific) following kit instructions. Genes were amplified from plasmids using specific primers: T7-*caspa* mRNA F: 5’-taatacgactcactatagggagagaatggccaaatctatcaagga-3’, *caspa* mRNA R: 5’-tcagagtccggggaacagg-3’, T7-*caspb* mRNA F: 5’-taatacgactcactatagggagaatggaggatattacccagct-3’, *caspb* mRNA R: 5’-tcacagtccaggaaacaggta-3’, T7-*Casp1* mRNA F: 5’-taatacgactcactatagggagaatggctgacaagatcctgag-3’, *Casp1* mRNA R: 5’-ttaatgtcccgggaagaggt-3’, T7-*Casp11* mRNA F: 5’-taatacgactcactatagggagaatggctgaaaacaaacaccct-3’, *Casp11* mRNA R: 5’-tgagttgccaggaaagaggt-3’. For overexpression experiments, 150 pg of mRNA were microinjected into the cell of a 1-cell stage zebrafish egg.

### Western Blotting

Cells were lysed in 1x RIPA buffer (Cell Signaling) with protease inhibitors (Roche), resolved on a 12% SDS-PAGE precast gel (BioRad) and transferred onto a 0.2 um PVDF membrane (BioRad). The membrane was blocked with 5% v/v milk in TBS buffer at room temperature for 2 h. The immunodetection was conducted by incubating the membrane with the anti-Casp11 (1:1000, Cat#180673, Abcam) or anti-Actin (1:1000, Cat#4968, Cell Signaling Technologies) antibodies for 12 h at 4° C under slow stirring. The membrane was then incubated with the secondary antibody conjugated to horseradish peroxidase (1:1000) for 2 h at RT. Target proteins were visualized by chemiluminiscence using the Clarity Western ECL Substrate (BioRad).

### Sytox Green cell uptake

Impermeable Sytox Green Nucleic Acid Stain (ThermoFisher Scientific) was used following manufacturer instructions at a final concentration of 5 μM to quantify cell death in RAW264.7 macrophages.

### FLICA staining

FAM-YVAD-FMK or 660-YVAD-FMK FLICA Caspase-1 Assay Kits (ImmunoChemistry Technologies) were used for zebrafish Caspa detection. The probe was diluted in egg water (1:200) and larvae were incubated during 1’5 hours at 28°C. Larvae were profusely washed in egg water before mounting and *in vivo* confocal imaging.

### *Mmar* culture DNA staining

Bacterial cultures were incubated O/N in a 5 μM solution of Draq5 Fluorescent Probe Solution (ThermoFisher Scientific). Bacterial suspensions were washed 3 times with PBS before RAW264.7 macrophages were infected.

### TUNEL assay

PFA fixed zebrafish larvae were used in combination with the *In situ* Cell Death Detection Kit (TMR and Fluorescein) (Roche). After O/N fixation larvae were washed and permeabilized for 45 min at 37°C using Proteinase K. TUNEL reactions were performed at 37°C O/N following kit instructions.

### Caspase activity assay

Caspase activity was assayed with the fluorometric substrate Z-YVAD-AFC (Santa Cruz) using 40 larvae per biological replicate as described previously (77). The fluorescence was measured in an Infinite M1000 microplate reader (Tecan) at an excitation wavelength of 400 nm and an emission wavelength of 505 nm.

### LysoTracker staining

Zebrafish larvae were immersed in egg water containing 10 μM LysoTracker Red DND-99 solution (ThermoFisher Scientific) for 1 hour. Before mounting and imaging larvae were washed 3 times with egg water.

### Lentiviral shRNA generation and validation

shRNA’s were acquired from the Mission library (Sigma, Zwijndrecht, the Netherlands) and were kindly provided by (dept. of Virus and Stem Cell Biology, LUMC, Leiden). For each gene 3 clones were selected for subsequent selection of the 2 most potent knockdowns. For *Casp11* knockdown TRCN0000012269 and TRCN0000012270 (Sigma-Aldrich) were selected. For *Gsdmd* knockdown TRCN0000219620 and TRCN0000198776 (Sigma-Aldrich) were finally selected.

HEK293T cells (ATCC, USA) were used for the generation of lentiviral particles as described in Heitzer et al. (78) using virulence and packaging vectors (pMD2.G (Addgene no. #12259) and psPAX2 (Addgene no. #12260)) kindly provided by Didier Trono (unpublished) with final concentrations of 0,72 pmol, 1,3pmol and 1,64pmol for virulence, packaging and shRNA plasmids respectively. Lentiviral supernatant was aliquoted and at −80°C prior to use. Lentiviral titer was determined using the Lentivirus-Associated p24 ELISA Kit (Cell biolabs) following suppliers instructions.

RAW 264.7 macrophage cultures (ATCC) were maintained in DMEM-F12 containing 10% FCS, and were transduced with 4 multiplicity of infection (MOI) scrambled control shRNA containing lentiviral particles or 4MOI of targeting shRNA containing viral particles in medium supplemented with 8µg/ml DEAE-dextran (Sigma). Medium was exchanged after 24 hours and puromycin (Gibco) selection (1,5µg/ml) was started 48 hours post transfection. After two passages (one week) under puromycin selection the remaining resistant clones were assessed for KD efficacy by qPCR using the following primers: qPCR *mmGsdmd* F: 5’-catcggcctttgagaaagtg-3’, qPCR *mmGsdmd* R: 5’-tcctgttcagaaggcagtag-3’, qPCR *mmCasp11* F: 5’-tggaagctgatgctgtcaag-3’, qPCR *mmCasp11* R: 5’-agcctcctgttttgtctgc-3’, qPCR *mmGapdh* F: 5’-atggtgaaggtcggtgtga-3’, qPCR *mmGapdh* R: 5’-ctggaacatgtagaccatgt-3’.

### Microscopy

Zebrafish larvae were imaged using a MZ16FA stereo fluorescence microscopy with DFC420C camera (Leica). Confocal z-tacks of zebrafish larvae and cell cultures were obtained on a TCS SP8 (Leica) using a 40x oil-immersion objective (NA 1.30). Before confocal imaging larvae were mounted in 1,5% low melting agarose in egg water containing tricaine as anesthetic. Images were processed and quantified using QuantiFish (http://doi.org/10.5281/zenodo.1182791) for stereo images or Fiji (79) for confocal images.

### Statistical Analysis

Statistical analysis was performed using GraphPad Prism 6 program (GraphPad Software). Comparisons between two groups were done using Mann-Whitney test. For analyzing differences between multiple groups ordinary one-way ANOVA + Tukey’s multiple comparisons test was used. Statistical significance was defined as *p<0.05, **p<0.01, ***p<0.001, ****p<0.0001. Graphs were plotted using PlotsofData (https://github.com/JoachimGoedhart/PlotsOfData).

## ACKNOWLEDGMENTS

We thank other members of the Meijer Laboratory for helpful discussions. We also thank the helpful fish facility and microscope facility staff at the Institute of Biology Leiden. We thank Dr. Paola Kuri and Dr. Maria Leptin for sharing the *asc:asc-EGFP* zebrafish line, Dr. George Lutfalla for the *mpeg1:mCherry* zebrafish line, Dr. Sander van Kasteren for kindly providing RAW264.7 macrophages, and Martijn Rablink for kindly providing shRNA plasmids. The work was funded by grant H2020-MSCA-IF-2014-655424 (M.V) from the European H2020 Marie Skłodowska-Curie Intra-European Fellowship program and by grant 13259 (A.H.M.) from the Netherlands Organization for Scientific Research, Domain Applied and Engineering Sciences (NWO-TTW).

## AUTHOR CONTRIBUTIONS

Conceptualization M.V and A.H.M.; Formal Analysis M.V. and A.H.M.; Investigation M.V.; Resources M.vd.V., A.G.; Writing – Original Draft M.V. and A.H.M.; Writing – Review & Editing M.V., M.vd.V., A.G. and A.H.M.; Visualization M.V.; Funding Acquisition M.V. and A.H.M.

## DECLARATION OF INTERESTS

The authors declare no competing interests.

## SUPPLEMENTARY FIGURES

**Supplementary Figure 1 (related to Figure 1).**
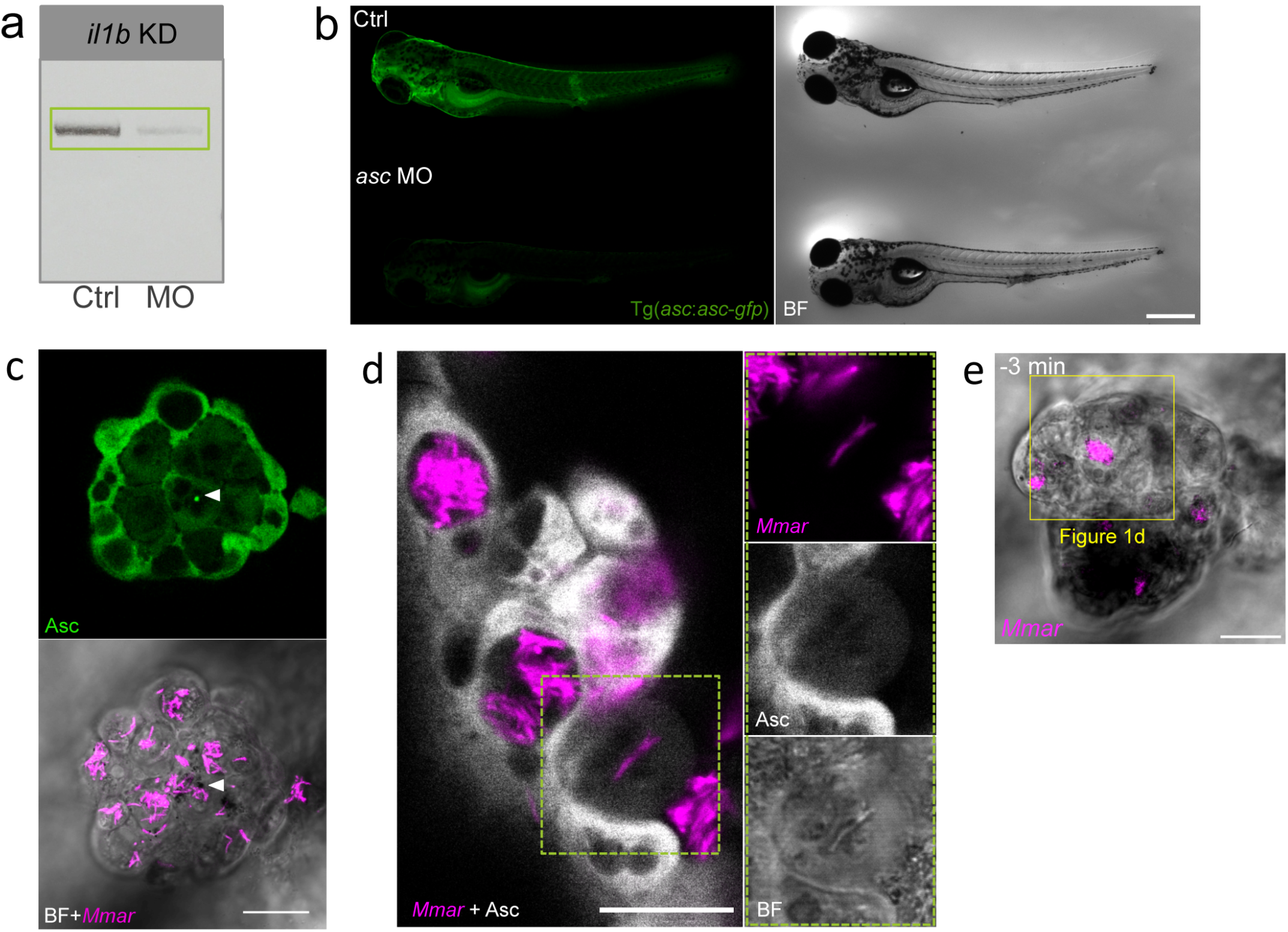
Asc-independent cell death is commonly visible in zebrafish granulomas. a, *il1b* KD efficiency was assessed by PCR on 4 days post-fertilization (dpf) zebrafish DNA samples. b, *asc* KD efficiency was assessed injecting *asc* MO into 1 cell-stage eggs from tg(*Asc:asc-EGFP*) zebrafish. Loss of Asc-GFP was assessed at 4 dpf. c, Confocal images of a zebrafish granuloma at 3 days post-infection (dpi) showing 1 Asc speck (arrowhead). d, Confocal and BF images of Asc speck-independent cell death in a zebrafish granuloma at 3dpi. e, Confocal image of the granuloma shown in Figure 1d. Scale bars are 500 (b) and 20 (c, d, e) μm.

**Supplementary Figure 2 (related to Figure 2).**
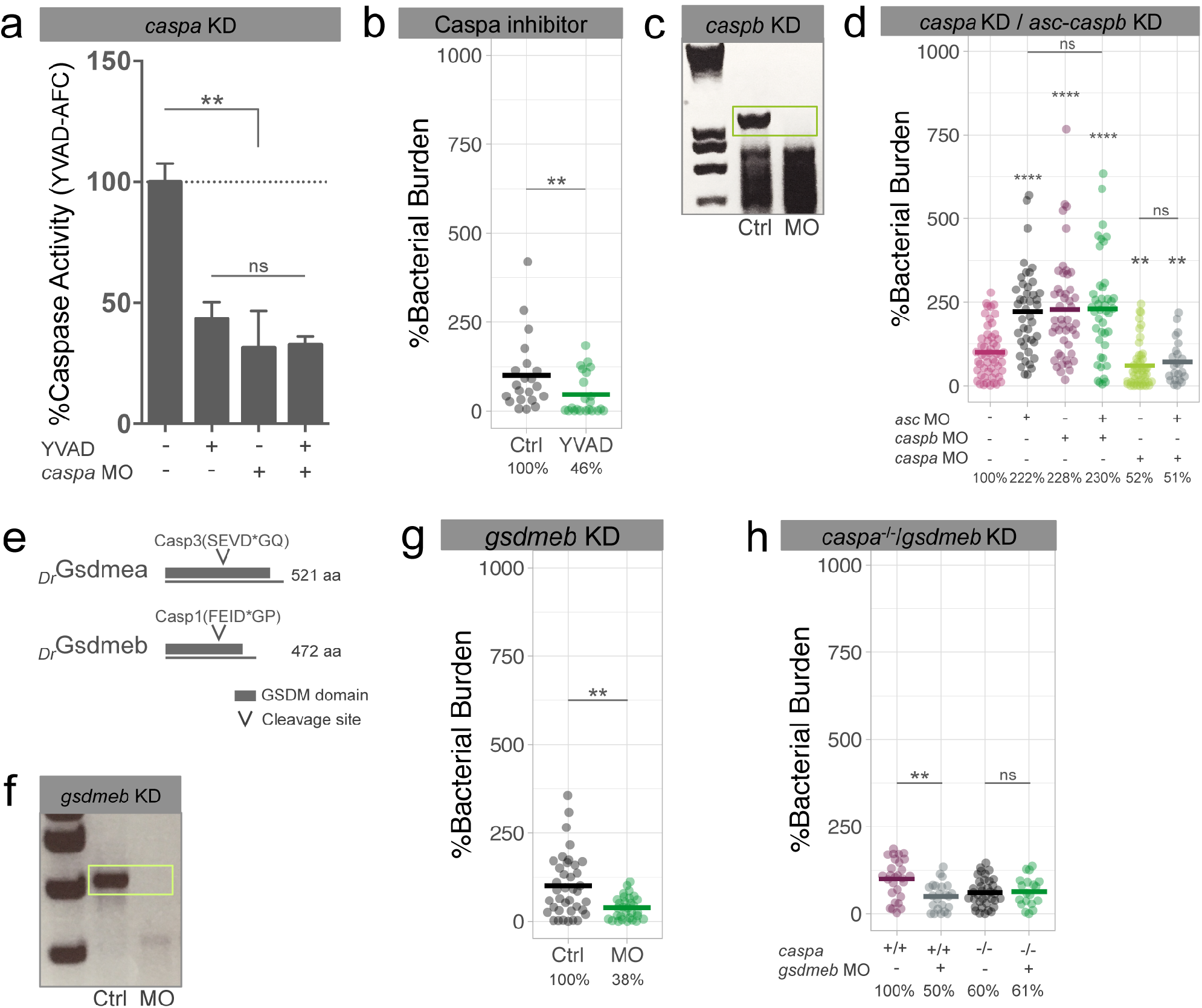
Disrupting the Caspa/Gsdmeb axis increases host resistance to *Mmar* infection. a, Caspase activity (YVAD-AFC) in control and *caspa* KD zebrafish showing that YVAD specifically inhibits Caspa in this infection model. b, Bacterial burden quantification after treatment with Caspa inhibitor (YVAD). c, *caspb* KD efficiency was assessed by PCR on 4 days post-fertilization (dpf) zebrafish DNA samples. d, Bacterial burden quantification in *caspa*+*asc* KD and *asc+caspb* KD. e, Schematic representing the 2 proteins with gasdermin domain in zebrafish (Gsdmea and Gsdmeb) and their caspase cleavage predicted sites. f, *gsdmeb* KD efficiency was assessed by PCR on 4 dpf zebrafish DNA samples. g, Bacterial burden data after transient *gsdmeb* KD in zebrafish larvae at 2 days post-infection (dpi) (300 cfu). h, Bacterial burden quantification in *caspa* mutants combined with *gsdmeb* KD at 2 dpi (300 cfu). Mann-Whitney test (b, e) and Ordinary one-way ANOVA + Tukey’s multiple comparisons test (a, d, h), **p<0.01, ****p<0.0001.

**Supplementary Figure 3 (related to Figure 3).**
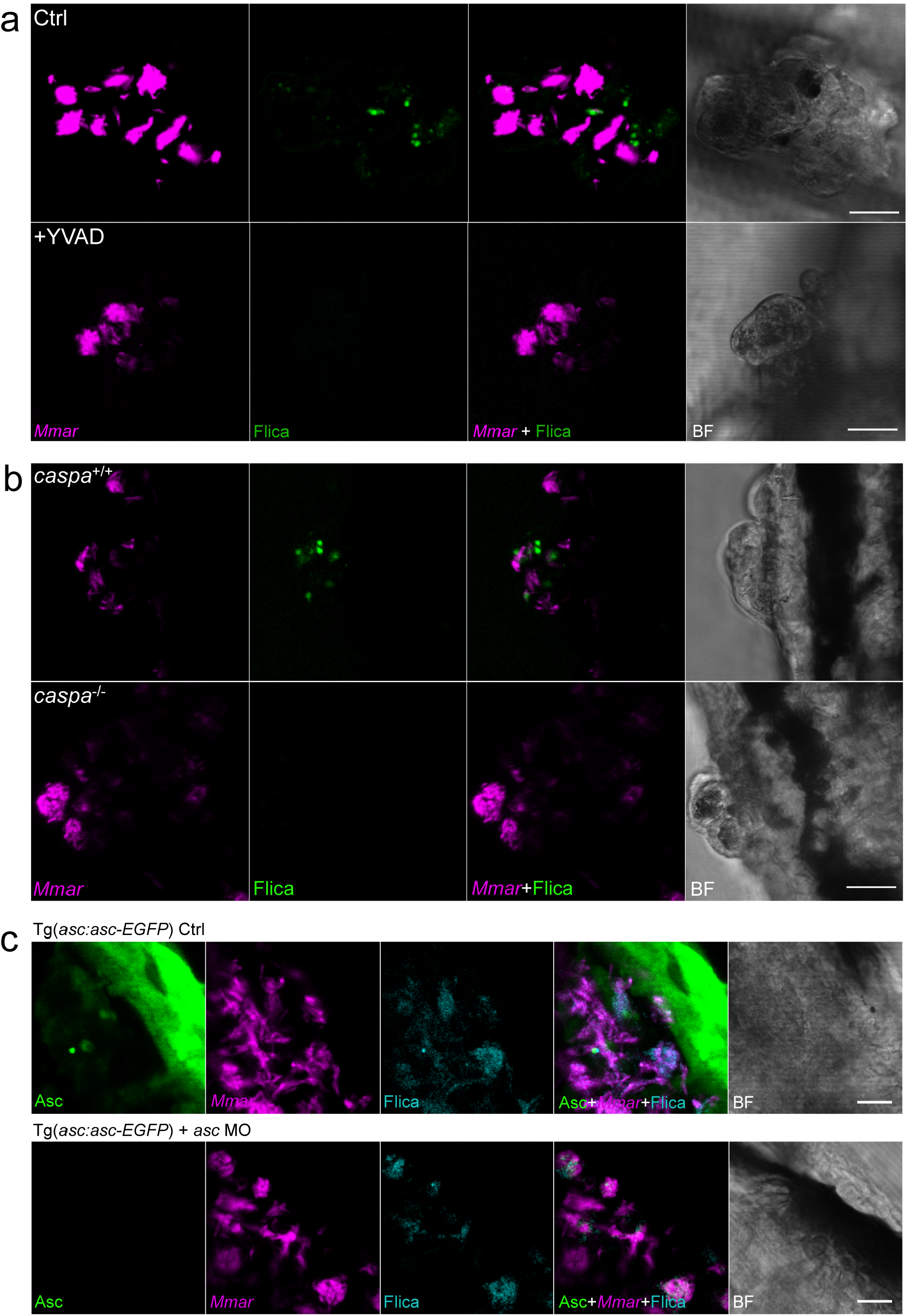
Asc-independent Caspa activation is specifically detected in zebrafish granulomas. a, Confocal images showing the specificity of Flica staining after treatment with YVAD. b, Confocal images showing the lack of Flica-positive signal in *caspa*^−/−^ zebrafish larvae. c, Confocal images showing Asc-independent Caspa activation (Flica) in *asc* KD *Mmar*-infected *asc:asc-EGFP* zebrafish larvae. Images are representative of >3 granulomas per condition. Scale bars are 20 (a, b, c) μm.

**Supplementary Figure 4 (related to Figure 5).**
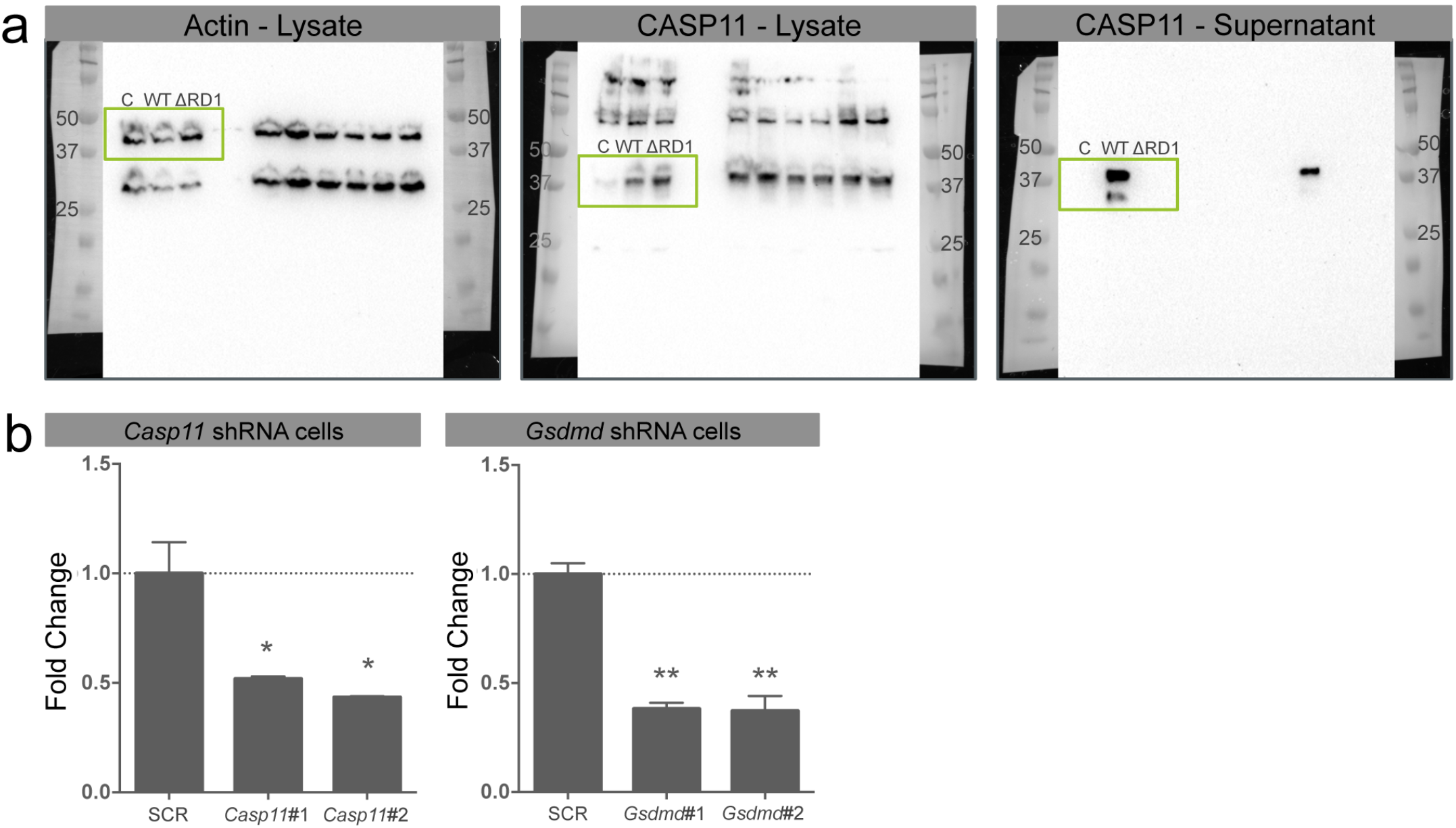
Silencing of CASP11 and GSDMD in RAW264.7 macrophages supports the requirement of these proteins for *Mmar*-induced pyroptosis. a, Full blots from Figure 5a. b, shRNA efficiency was assessed by qPCR on RAW264.7 DNA samples. Mann-Whitney test (b), *p<0.05, **p<0.01.

**Supplementary Figure 5 (related to Figure 7).**
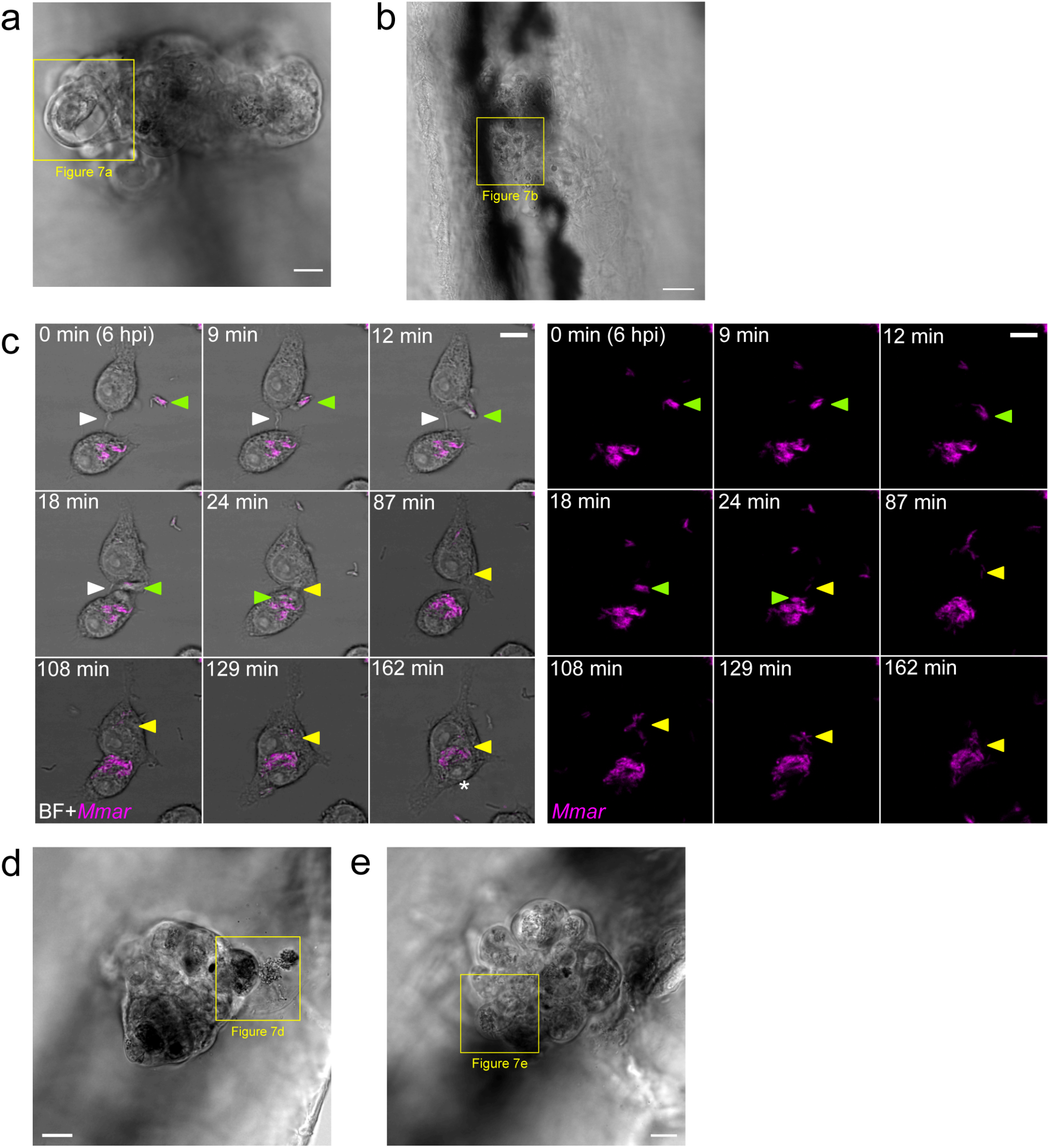
*Mmar* infection induces formation of cell-in-cell structures *in vivo* and *in vitro*. a, Zebrafish granuloma in Figure 7A. b, Zebrafish granuloma in Figure 7B. c, Confocal time-series showing cell-in-cell formation during *Mmar* infection (MOI=10) in RAW264.7 macrophages, shown with BF (left) or fluorescence only (right). Depicted cells are in contact (white arrowhead). *Mmar* cluster (green arrowhead) appears. Cell 1 (top cell) takes it up and transfers it inside cell 2 (bottom cell). Some bacteria from the cluster remain inside cell 1 and migrate towards the other cluster at the top of the cell (yellow arrowhead) between min 24 and 87. Entosis starts in min 108. Bacteria inside the top cell migrate towards the bottom cell (yellow arrowhead) while this cell is internalized by the top cell between min 108 and 129. Asterisk in min 162 points out nuclear shrinkage in the inner cell. See also Supplementary Video 5. d, Zebrafish granuloma in Figure 7d. e, Zebrafish granuloma in Figure 7e. Scale bars are 10 (a, b, c, d, e) μm.

**Supplementary Figure 6 (related to Figure 7).**
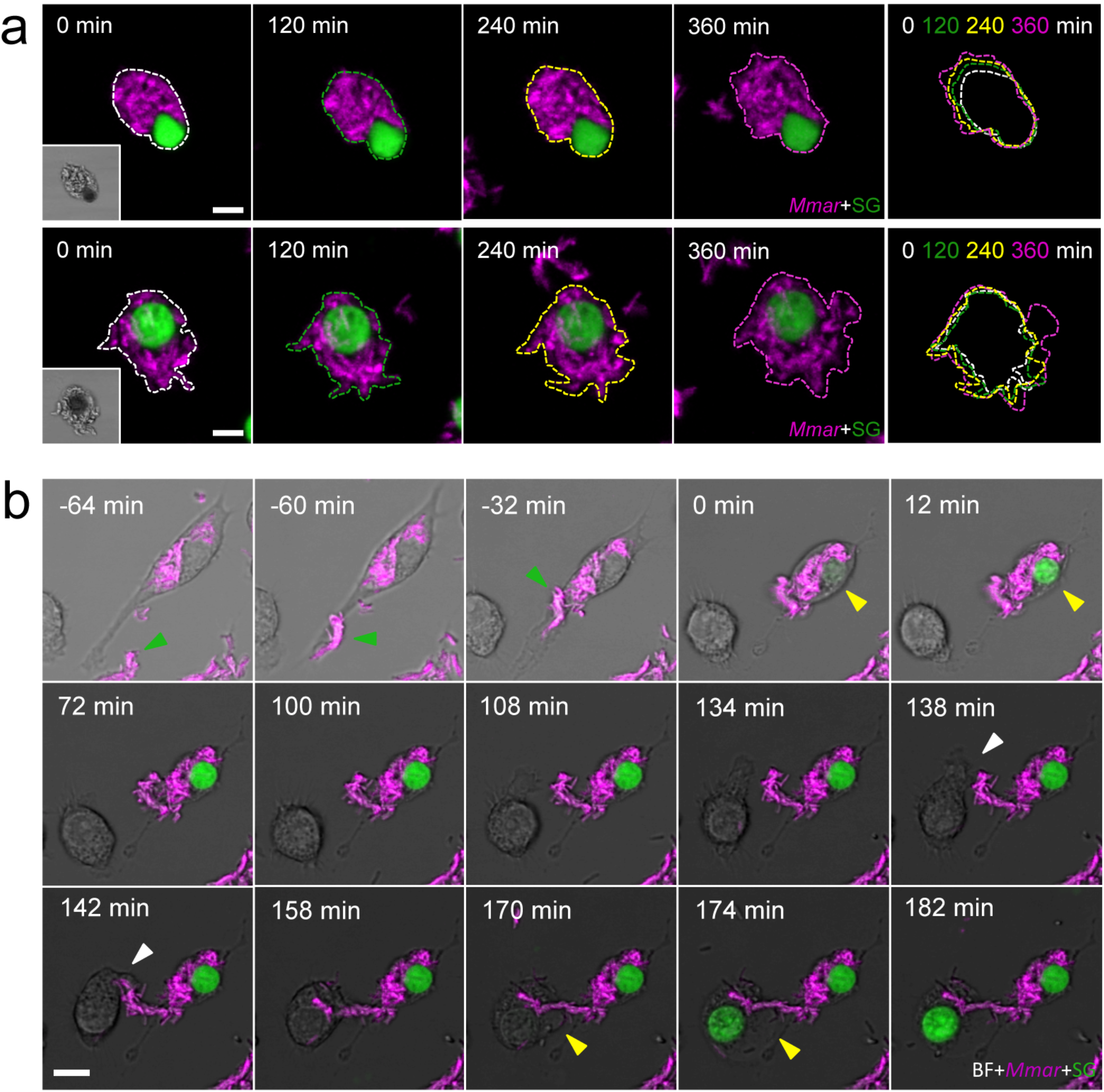
Pyroptosis of infected cells leads to *Mmar* extracellular growth and infection amplification. a, Two examples of extracellular growth of *Mmar* after RAW264.7 cell pyroptosis during 360 min after cell death. Expansion of extracellular growth is indicated by the overlay of coloured outlines at different time points. b, A previously infected macrophage phagocytoses a new *Mmar* cluster (green arrowhead) and dies via pyroptosis (yellow arrowhead). The dead macrophage attracts a new macrophage to the area. The newly arrived macrophage phagocytoses *Mmar* (white arrowhead) and dies via pyroptosis (yellow arrowhead) (see also Supplementary Video 6). Scale bars are 10 (a, b) μm.

## SUPPLEMENTARY VIDEOS

**Video 1 (related to Figure 1d). Asc speck is degraded after pyroptosis and phagocytosis by a new cell in a zebrafish *Mmar* granuloma.** The video shows two channels, BF+Mmar on the left and Asc (Fire LUTs) on the right. Confocal time-lapse shows Asc speck formation in Cell 1 (green line) and Asc speck degradation by Cell 2 (white line) after phagocytosis. Time in hh:mm.

**Video 2 (related to Figure 3d). *Mmar* triggers ASC-independent and phagosome rupture-dependent macrophage pyroptosis.** Confocal time-lapse show a *Mmar* cluster being phagocytosed by a macrophage (white arrowhead). Phagosome rupture (pink arrowhead) and subsequent cell pyroptosis (yellow arrowhead) are visible minutes after phagosome formation. Time in hh:mm.

**Video 3 (related to Supplementary Figure 5c). *Mmar* infection induces formation of cell-in-cell structures *in vitro.*** Confocal time-series showing cell-in-cell formation during *Mmar* infection in RAW264.7 macrophages. White arrowheads point cell-cell contact occurring before cell-in-cell formation. The top cell takes a *Mmar* cluster (green arrowhead) and transfer it to the bottom cell. Some bacteria remain inside the top cell and migrate towards the top of the cell (yellow arrowhead). Bacteria inside the top cell migrate towards the bottom cell (yellow arrowhead) while this cell is internalized by the top cell. Time in hh:mm.

**Video 4 (related to Figure 7d). Pyroptosis initiation induces the release of the inner cell from the cell-in-cell structure.** The video shows two channels, BF+M*mar* on the left and Asc (inverted LUTs) on the right. Confocal time-lapse showing the release of the inner cell from the cell-in-cell structure following the initiation of pyroptotic cell death (green arrowheads). The infected outer cell eventually dies via pyroptosis (yellow arrowheads) 1 hour later. Time in hh:mm.

**Video 5 (related to Figure 7e). Cell-in-cell structures in granulomas can be formed after sequential uptake of multiple infected cells.** The video shows two channels, BF+*Mmar* on the left and Asc (inverted LUTs) on the right. Confocal time-lapse showing how a single cell in a zebrafish granuloma can sequentially take up more than 1 infected cell (Cell 1 in yellow, Cell 2 in green, Cell 3 in white). Time in hh:mm.

**Video 6 (related to Supplementary Figure 6b). Pyroptosis of infected cells leads to *Mmar* infection amplification.** Confocal time-lapse showing a previously infected macrophage phagocytosing a new *Mmar* cluster (green arrowhead) and dying via pyroptosis (SytoxGreen uptake). Dead macrophage attracts a new macrophage to the area. Newly arrived macrophage phagocytoses *Mmar* (white arrowhead) and quickly dies via pyroptosis (yellow arrowhead). Time in hh:mm.

